# Molecular dissection of zinc-mediated immunity in Arabidopsis thaliana

**DOI:** 10.64898/2026.08.28.747889

**Authors:** Viviana Escudero, Cuong V. Hoang, Antoni Garcia-Molina, Aishee De, Alejandro M. Armas, Dennis Brueckner, Darío Ferreira Sánchez, Christoph Bueschl, Maria Doppler, Antony van de Ent, Rainer Schuhmacher, Manuel González-Guerrero, Lucía Jordá

**Author notes:** **Correspondence and material distribution:** Lucía Jordá, Manuel González-Guerrero. Centro de Biotecnología y Genómica de Plantas, Universidad Politécnica de Madrid, Instituto Nacional de Investigación y Tecnología Agraria y Alimentaria/Consejo Superior de Investigaciones Científicas, Pozuelo de Alarcón, 28223 Madrid, Spain. Dept. of Microbiology & Plant Pathology, University of California, Riverside, CA, 92521.

## Abstract

Zinc is an essential micronutrient at low concentrations, yet it becomes toxic at slightly higher ones. This is exploited by plants as an effective defensive strategy. However, the molecular components that are involved zinc-mediated immunity remain poorly defined. Here, we show that mixed-linked β-1,3/1,4-glucans naturally ocurring in microbial and grass cell walls and used as an agrobiological solution, trigger zinc accumulation in the Arabidopsis apoplast and upregulate the expression of the zinc transporters HMA2 and HMA4. This response occurs independently of salicylic acid, jasmonic acid and ethylene-mediated signalling pathways, but it requires the LysM receptor kinases CERK1, LYK4 and LYK5, indicating a specific pattern triggered immunity-associated mechanism. We further demonstrate that *hma2hma4* mutants display constitutive activation of a broad set of defence-related genes, yet this transcriptional reprogramming is insufficient to confer resistance against the necrotrophic fungus *Plectosphaerella cucumerina* BMM. Moreover, metabolomic profiling highlights the contribution of specialized metabolites to this defective defence output. Altogether, our findings reveal that zinc-mediated toxicity constitutes a defence mechanism integrated into the immune response triggered by specific microbial or damage associated molecular patterns.

## 1. Introduction

Transition elements such as iron, copper, or zinc are essential nutrients at low concentrations, as they are key cofactors of enzymes participating in almost all biological processes (Fraustro da Silva and Williams, 2001). However, these elements become toxic at slightly higher levels, since they can produce free radicals in non-enzymatic reactions or displace the native metal cofactor in some metalloenzymes (Goldstein et al., 1993; Kramer, 2024; Macomber and Imlay, 2009). Growing evidence indicates that plants use this double-edged nature of transition metal biology to combat pathogens (Cabot et al., 2019; Dangol et al., 2019; De et al., 2024; Morina and Küpper, 2022). As it also occurs in animals, where nutritional immunity limits pathogen proliferation by restricting access to essential micronutrients (Murdoch and Skaar, 2022), plants remove key metals from infection sites as part of their defence strategy (Cao et al., 2024). Alternatively, plants can adopt an opposite strategy and locally increase metal concentrations to levels that become toxic for the invader. This is further reinforced by the accumulation of apoplastic metalloenzymes involved in plant immunity (Escudero et al., 2022; Morina et al., 2021).

Zinc-mediated toxicity to combat infection is widely extended, having been observed in Arabidopsis, soybean, and pepper plants (Escudero *et al*., 2022; Kuvelja et al., 2024; Morina *et al*., 2021). This layer of the immune response relies on the accumulation of zinc at the infection site, a process facilitated by Zn^2+^-transporting ATPases such as *Arabidopsis thaliana* HMA2 and HMA4 (Escudero *et al*., 2022). Consequently, optimal zinc nutrition may enhance plant pathogen resistance (Kuvelja *et al*., 2024). In response to host-induced zinc-toxicity, pathogens activate their own metal-detoxification mechanisms, which can confer them with a competitive advantage during host-colonization (Visconti et al., 2022). Beyond the roles of HMA2 and HMA4, and the observed dysregulation of marker genes associated with salicylic acid (SA), jasmonic acid (JA) and ethylene (ET) signalling pathways (Escudero *et al*., 2022), little is known about the molecular players that trigger zinc-mediated immunity (ZiMI), how this signal is transduced, the sources of mobilized zinc, or how plant physiology accommodates for this response.

To counter pathogen threats, plants rely on a robust innate immune system that senses both conserved non-self molecules from microorganisms (Microbe-Associated Molecular Patterns, MAMPs) and self-derived molecules released upon tissue damage (Damage-Associated Molecular Patterns, DAMPs) (Molina et al., 2024b). Recognition of these signals triggers a cascade of defence responses that preserve cellular integrity and plant fitness (DeFalco and Zipfel, 2021). This cell-intrinsic surveillance system is mediated by a highly complex network of plasma membrane-anchored pattern-recognition receptors (PRRs), which initiate a strong defence response known as Pattern-Triggered Immunity (PTI) upon MAMP/DAMP recognition through their extracellular domains (ECDs) (Snoeck et al., 2025; Yu et al., 2017). Proteins and carbohydrates constitute major structural components of both host and microbial surfaces and act as pivotal signals (MAMPs and DAMPs) during the early stages of plant-pathogen interactions. Representative MAMPs include the bacterial peptides flg22 and elf18, as well as glycans with different degrees of polymerization derived from fungal and oomycete walls, such as chitin oligomers (e.g., chitohexaose, CHI6), unbranched mixed-linked β-1,3/1,4-D-glucans (MLGs), or linear β-1,2-D-glucans and peptidoglycans from bacteria (Chinchilla et al., 2007; Fuertes-Rabanal et al., 2024; Kunze et al., 2004; Liu et al., 2012; Rebaque et al., 2021; Willmann et al., 2011). DAMPs comprise peptides released upon pathogen attack, such as *Arabidopsis thaliana* PLANT ELICITOR PEPTIDE 1, *At*PEP1 or SERINE RICH ENDOGENOUS PEPTIDES, SCOOPs (Xiao et al., 2025), and a wide array of oligosaccharides of different lengths that originate from plant cell wall polysaccharides through the action of cell wall degrading enzymes (CWDEs) secreted by pathogens (Molina et al., 2024a). Examples include cello-oligomers (e.g. cellotriose, CEL3), MLGs from grasses, and arabinoxylans (e.g., 3^3^-α-L-arabinofuranosyl-xylotetraose, XA^3^XX), among other wall-derived carbohydrates (Sun et al., 2025), whose capacity to trigger early immune responses has allowed their use as agrobiological solutions for sustainable agriculture (Rebaque et al., 2021). The Arabidopsis genome comprises more than 600 genes encoding membrane-bound receptors (Liu et al., 2024), which are classified according to the structure of their ECDs with leucine-rich repeat (LRR) and lysin motif (LysM) domains being among the best characterized extracellular recognition domains; this ECD architecture determines the type of ligand they recognize. Binding of specific DAMPs or MAMPs to the PRR ECD activates PTI, leading to transient cytoplasmic Ca^+2^ influxes, phosphorylation cascades involving mitogen-activated protein kinases (MAPKs), and the production of reactive oxygen species (ROS). These early signalling events trigger large transcriptional reprogramming, resulting in the synthesis of defence-related phytohormones, antimicrobial peptides, and specialized metabolites, alongside metabolic alterations to limit pathogen access to key nutrients and promote the accumulation of antimicrobial compounds (Campos et al., 2018; Couto and Zipfel, 2016; Singh et al., 2023).

In this work, we have determined that ZiMI constitutes a defence mechanism activated during PTI. We show that active zinc pumping into the apoplast occurs after the perception of carbohydrate-based MAMPs or DAMPs, such as MLG43. Mutants defective in receptors involved in MLG perception exhibit impaired apoplastic zinc accumulation in response to this carbohydrate-based signalling molecule. A ZiMI-related transcriptional program has been identified through comparative transcriptome analysis of wild type and *hma2hma4* plants under mock conditions and following *Plectosphaerella cucumerina* BMM (*Pc*BMM) infection. This transcriptional reprograming leads to profound changes in the metabolome, including the loss of synthesis of known antimicrobial compounds and altered expression of genes related to glucosinolate and phenylpropanoid metabolism.

## 2. Material and Methods

### 2.1. Plant materials and growth conditions

*Arabidopsis thaliana* plants from Col-0 ecotype were used in this study, including the following mutants, *hma2hma4* (Hussain et al., 2004), *cerk1-2lyk4-1lyk5-1, igp1-6* and *igp4-1* (Martín-Dacal et al., 2023). Plants for RNAseq, metabolomic analysis, X-Ray fluorescence and pathogen inoculations, were grown in a mixture of peat:vermiculite (3:1), stratified at 4 ⁰C and moved to a growth chamber under short day conditions (10 h light photoperiod, ∼150 µEm^-2^s^-1^), 65% humidity and 20-22 ⁰C. For gene expression analyses 12 to 15 Col-0 and mutant seeds were grown in 24-well plates under long day conditions for 14 days (16 h light photoperiod, ∼150 µEm^-2^s^-^) at 20-22 ⁰C in liquid ½ MS medium with 1 % saccharose.

### 2.2. Pathogen inoculations

Pathogen inoculations using the necrotrophic fungus *Plectosphaerella cucumerina* BMM (*Pc*BMM) were performed as described in Escudero et al., 2022.

### 2.3. Carbohydrates and peptides used in the experiments

The following oligosaccharides, hexaacetyl-chitohexaose (CHI6; β-1,4-D-(GlcNAc)_6_; O-CHI6; Sigma), cellotriose (CEL3; β-1,4-D-(Glc)_3_; O-CTR; Megazyme), 3^3^-α-L-arabinofuranosyl-xylotetraose (XA^3^XX; O-XA3XX: Megazyme), and unbranched mixed-linked glucan (MLG43; 3^1^-β -D-cellobiosyl-glucose; 0-BGTRIB; Megazyme) were diluted in sterile water and used at the final concentrations indicated in the assays. The peptidic MAMP used in this study, flg22 (QRLSTGSRINSAKDDAAGLQIA) was synthesized by Genscript.

### 2.4. Synchrotron-based X-ray Fluorescence assays

Leaves from 16-17-day-old Arabidopsis plants grown on soil were drop-inoculated with a 2.5 µl droplet of *Pc*BMM spore suspension (4 x 10^6^ spores/ml^-1^), covered with a plastic foil, and collected at 48 hpi. Leaves were detached and mounted in a tight sandwich between two sheets of Ultralene membrane. Synchrotron radiation scanning micro-XRF was performed at the Swiss Light Source (SLS; microXAS beamline, Villigen, Switzerland) following conditions described in Escudero et al., 2022. For DAMP/MAMPs treatments, 5 µl drops were applied to leaves from 16-17-day-old Arabidopsis plants, using the following concentrations: 1 µM flg22, 50 µM CHI6, 10 µM CEL3, 450 µM XA^3^XX, and 50 µM MLG43, covered for 20 h with a plastic foil, and collected 24 h post-treatment. Leaves were mounted between two sheets of Ultralene membrane. The spatial distribution of Zn and Ca was performed by synchrotron-based X-ray fluorescence (S-XRF) scanning at the Deutsches Elektronen-Synchrotron, DESY (PETRA III, Beamline P06). An excitation energy of 12 keV was used to acquire the images shown in Figures 1A and D. X-ray fluorescence (XRF) signals were collected using two 4-element Vortex ME4 silicon drift detectors (SDDs; Hitachi High Technologies America, Inc.), positioned at 45° and 315° relatively to the forward beam direction. Prefocusing compound refractive lenses (CRLs) were used to increase the photon flux at the expense of a larger beam size, resulting in a focused beam size of 2.9 µm x 1.9 µm (horizontal x vertical). The beam was focused with a photon flux at focus of 1.5x 10^11^ ph/s. For the images shown in Figure 1B, the primary X-ray energy was set to 16keV. XRF signals were collected using a 16-element ASCANIO SDD (Politecnico di Milano), positioned at 180° relative to the forward beam direction in backscatter geometry. The focused beam size was 460nm x 440nm (horizontal x vertical) with a photon flux at focus of 7.6x 10^9^ ph/s.

**Fig. 1.**
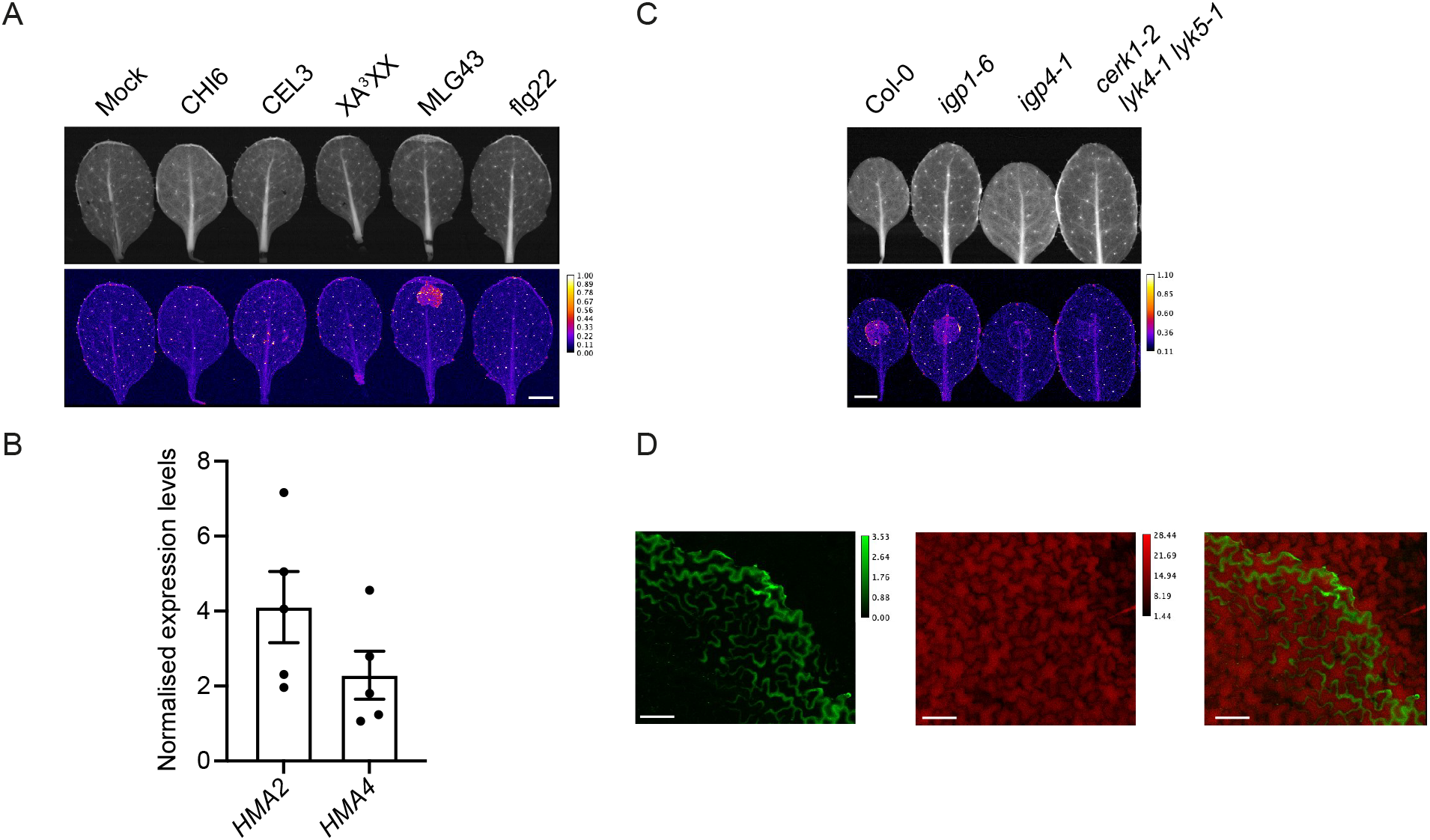
Mixed-linked glucan oligosaccharide MLG43 triggers strong ZiMI. (A) Synchrotron-based X-ray fluorescence (S-XRF) images of zinc accumulation in Col-0 plants 24 hours post 5 µl drop-application of distilled water (mock), 50 µM CHI6, 10µM CEL3, 500 µM XA^3^XX, 50 µM MLG43, and 1 µM flg22. Units indicate areal density µg/cm2. Each image is representative of three independent replicates showing similar results. Scale bar = 2 cm. Upper panel shows transmission map. (B) Relative expression levels of *HMA2* and *HMA4* at 6 h upon 50 µM MLG43 application. Expression levels relative to the *UBC21* gene (At5g25760) are shown. Data represent the mean of three technical measurements from three biological replicates. (C) S-XRF images of zinc in wild type Col-0, *igp1-6, igp4-1*and *cerk1-2lyk4-1lyk5-1* mutant plants 24 hours post treatment with 50 µM MLG43. Units indicate areal density µg/cm^2^. Scale bar = 2 cm. Upper panel shows transmission map. (D) A higher-magnification view of the leaf area treated with 50 µM MLG43. Left panel shows the apoplastic zinc distribution in green, while potassium content is shown in red in the middle panel. Units indicate areal density µg/cm^2^. The right image shows the overlay of the two images, where no co-localization of zinc and potassium takes place in the MLG43-treated area.

### 2.5. Gene expression analyses

For RNAseq, 16-day-old Arabidopsis Col-0 and *hma2hma4* plants were mock or *Pc*BMM spray-inoculated, and tissues were harvested at 48 hpi. Each sample represented a pool of the rosettes of 10-15 plants grown under the growth conditions described. Total RNA, from three biological replicates, was extracted using a phenol-chloroform method. RNA-Seq libraries were prepared according to Illumina protocols and sequenced for 150 pair-end reads in an Illumina HiSeq2500 by Novogene (Cambridge, UK). RNA-Seq datasets were processed with a standard pipeline using Trimmomatic – RNA STAR-featureCounts – DESeq2 (Bolger et al., 2014; Dobin et al., 2013; Liao et al., 2014; Love et al., 2014) using the Arabidopsis genome annotation of TAIR10. Differentially expressed genes were selected according to absolute fold-changes ≥ 1 to respective controls (adjusted *P* ≤ 0.05, Wald’s test) or for the interaction of genotype and treatment (adjusted *P* ≤ 0.05, Wald’s test). GO categories statistically overrepresented (adjusted *P* ≤ 0.05, Fisher’s Exact test) in the differentially expressed set of genes were obtained with Metascape (Zhou et al., 2019). Hierarchical clustering heatmaps according to the Ward D2 method were drawn using z-scores of normalised counts of changing transcripts with the RStudio package *pheatmap*.

For gene expression analysis, 14-day-old Arabidopsis seedlings were grown on liquid ½ MS media and treated with DAMP/MAMPs (same concentrations used for Synchrotron-based X-ray Fluorescence assays) for 6 h. RNA was extracted using a phenol-chloroform method. DNase treatment (Invitrogen), cDNA synthesis (Roche) and qRT-PCR (SYBR, Applied biosystems) were performed following the manufacturer’s protocol. Oligonucleotides used are listed in Supplementary Table S10. Gene expression was normalized with the house keeping gene *UBC21* (At5g25760).

### 2.6. Metabolomic analysis

16-days old plants of Col-0 and *hma2hma4* plants were sprayed with mock (water) or with a *Pc*BMM spore suspension of 4×10^6^ spores/ml and covered with a plastic film. 48 h post inoculation, aerial tissues from 3 independent replicates were collected and Nitrogen frozen. Sample preparation and LC-HRMS analysis were performed as follows. Frozen plant material was weighed in (between 100 and 120mg) and milled directly in 2.0 mL tubes using a Retsch ball mill with respective liquid nitrogen pre-cooled adapters. Milled plant material was extracted with 10 volumes of extraction solvent [1:1.5:1.5 (v/v/v) H_2_O:acetonitrile:methanol] acidified with 0.1% formic acid. The samples were vortexed for 10 s, sonicated in an ultrasonic bath for 15 min, and centrifuged at 4°C (7500 rpm) for 10 min. For uniformly ^13^C-labeled material (Isolife, lyophilised, uniform, 97 atom% ^13^C, *Arabidopsis thaliana* Columbia-O) a typical water content of fresh leaf material was achieved by addition of the corresponding volume of water immediately before extraction. Aliquots of ^13^C-labeled extracts were added to each replicate of the native experimental samples at a 1:1 ratio /v/v). Subsequently, 100 µL of ^13^C extract and 100 µL of native extract were mixed with 100 µL H_2_O containing 0.1% formic acid, centrifuged at 4°C for 10 min at7500 rpm, and the supernatants were transferred to glass vials for liquid chromatography–high-resolution mass spectrometry (LC-HRMS) analysis. LC-HRMS measurements were carried out as described previously (Doppler et al., 2022); in brief, a QExactiveHFOrbitrap mass spectrometer coupled to a Vanquish HPLC system was operated in fast polarity-switching mode, and full-scan spectra were acquired in profile mode over an *m/z* range of 100–1000 at a resolving power setting of 120,000 (at *m/z* 200, FWHM (full width half maximum)). For LC-HRMS/MS, the data dependant mode with an inclusion list and separate positive and negative ionisation mode measurements were used. Inclusion lists consisted of the metabolite ion pairs (native and ^13^C form) detected in full scan mode. Product ion spectra were recorded with stepped collision energy (20 eV, 45 eV, 70 eV) with a resolving power setting of 30,000 (at *m/z* 200, FWHM).

Raw LC-HRMS data was converted to mzML format using ProteoWizard (version 3.0.23163; https://proteowizard.sourceforge.io/) and subsequently further processed with MetExtract II (Bueschl et al., 2017) for untargeted feature detection, relative quantification, and annotation. Parameters were Isotopic Enrichment 98.93 (12C) and 96% (13C), Intensity Threshold 1E3, Number of Isotopologs checked (12C and 13C) 2, Number of Carbon Atoms searched 3-60, Maximum allowed Isotopolog Ratio Error 35%, Maximum allowed MZ deviation (intra-scan) 3ppm, Extracted Ion Chromatogram width +/- 5 ppm, Chromatographic Width 3-19 scans, Minimum Correlation of Chromatographic Peaks for Convolution 0.85, Maximum Deviation for Bracketing of Features 10 ppm and 0.1 minutes. Detected features were used to generate inclusion lists for successive LC-HRMS/MS analysis (same instrument and LC method, up to 5 MSMS scans automatically triggered, resolving power of 30,000 at *m/z* 200 for MSMS scans). LC-HRMS/MS data was processed with MZmine (version 4.5.20, (Schmid et al., 2023). Workflow setup and parameters were (main steps) Spectral Library Import (MSMS spectra of in-house standards, MoNA Massbank (https://mona.fiehnlab.ucdavis.edu/), Japan Massbank (https://massbank.jp/MassBank/)), Mass Detection (Noise Level 1E4 for MS1 and 1E2 for MS2), Chromatogram Builder (M/Z Tolerance 10 ppm), Smoothing (Savitzky Golay 5), Local Minimum Feature resolver (Chromatographic Threshold 73.9%, Minimum Absolute Height 5E4, Peak Duration 0.1 – 3.0 minutes, Minimum Scans 3), Join Aligner (*m/z* Tolerance 8 ppm, Retention Time Tolerance 0.04 minutes), Spectral Library Search (Merged (simple) Spectra, Merging *m/z* Tolerance 25 ppm, Precursor Tolerance 5 ppm, Spectral *m/z* Tolerance 25 ppm, Minimum Matched Signals 4, Weighted Cosine Similarity, Minimum Cos. Similarity 0.7), Export to csv and mgf format. Subsequently, the MSMS spectra of the detected compounds were processed with SIRIUS (version 6.1.0, (Duhrkop et al., 2019) with settings set to Orbitrap. Identified compounds, annotated MSMS spectra, Database hits and SIRIUS/CANOPUS results were subsequently organized on a per feature level and restricted to hits where the number of carbon atoms matched the determined number of carbon atoms from the MetExtract II analysis (derived from the isotopic labeling experiment). Furthermore, putative chemical formulas were generated with the Seven Golden Rules (Kind and Fiehn, 2007). Annotations that are further discussed were manually reviewed.

Statistical analysis was implemented in R (version 3.5.3; https://www.r-project.org/). Missing values were replaced with 0 and the most abundant feature/adduct was selected for each metabolite (i.e., the one with the highest average peak area among all experimental samples). The critical alpha threshold was set to 0.05 and a minimum absolute log2 fold change of 1 was required for a feature to be designated to be statistically significantly different between the tested groups. For counting if a feature was present in a group or not it had to be present in at least 2 of the 3 replicates.

## 3. Results

### 3.1. Mixed linked glucan MLG43 elicits ZiMI response in A. thaliana

To determine the molecular pattern(s) triggering ZiMI, different MAMPs and DAMPs, such as chitohexaose (CHI6), cellotriose (CEL3), arabinoxylan (XA^3^XX), MLG43, and flagellin (flg22) were drop-applied to Arabidopsis leaves and zinc distribution was observed 24 hours-post-inoculation (hpi) using Synchrotron-based X-ray fluorescence (S-XRF) (Fig. 1A). Only CEL3 and MLG43 activated zinc accumulation in the treated leaves, eliciting MLG43 the strongest response. MLG43 also induced *HMA2* and *HMA4* expression in Col-0 plants (Fig. 1B), consistent with the role of these transporters in allocating Zn^2+^ for ZiMI (Escudero *et al.,* 2022). Similarly, loss of the proteins involved in MLG43 sensing should result in lower zinc-accumulation at the infection site. Although the receptor that binds MLG43 has not yet been identified in Arabidopsis, several PRRs have been shown to contribute to MLG43-triggered immune signalling (Martín-Dacal et al., 2023; Rebaque et al., 2021). Therefore, we tested whether mutants in the LysM receptor kinases (*cerk1-2lyk4-1lyk5-1* triple mutant) and in the *IGP* family (*igp1-6* and *igp4-1*) show altered zinc levels upon MLG43 treatment. While the *igp1 and igp4* single mutants displayed no detectable effect on the ZiMI response, the *cerk1-2lyk4-1lyk5-1* triple mutant presented a severe reduction of zinc accumulation in leaves (Fig. 1C). To confirm that this zinc was accumulating in the apoplast and that it was not caused by cell lyses, potassium was used as an indicator of the cell content as well as of cellular integrity. As it is shown in Fig. 1D, there was no co-localization of zinc and potassium in the MLG43-treated area, being zinc confined to the apoplastic space between the potassium-defined cells.

Zinc used during the immune response could conceivably originate either from the immediate vicinity to the infection site or be translocated from distal plant organs. To address this question, zinc distribution in mock-treated, MLG43-treated, and mechanically injured leaves were determined comparing samples detached from the plant prior to treatment with leaves that remained attached to the plant. As it can be observed in Fig. 2A, zinc accumulation upon MLG43 required that the leaf remained attached to the whole plant. The assay also showed that ZiMI is not due to tissue damage, supporting its position in the early immune response triggered by PTI. However, a functional root system was not required for ZiMI, as preserving rosette was sufficient to observe zinc accumulation when MLG43 was applied (Fig. 2B).

**Fig. 2.**
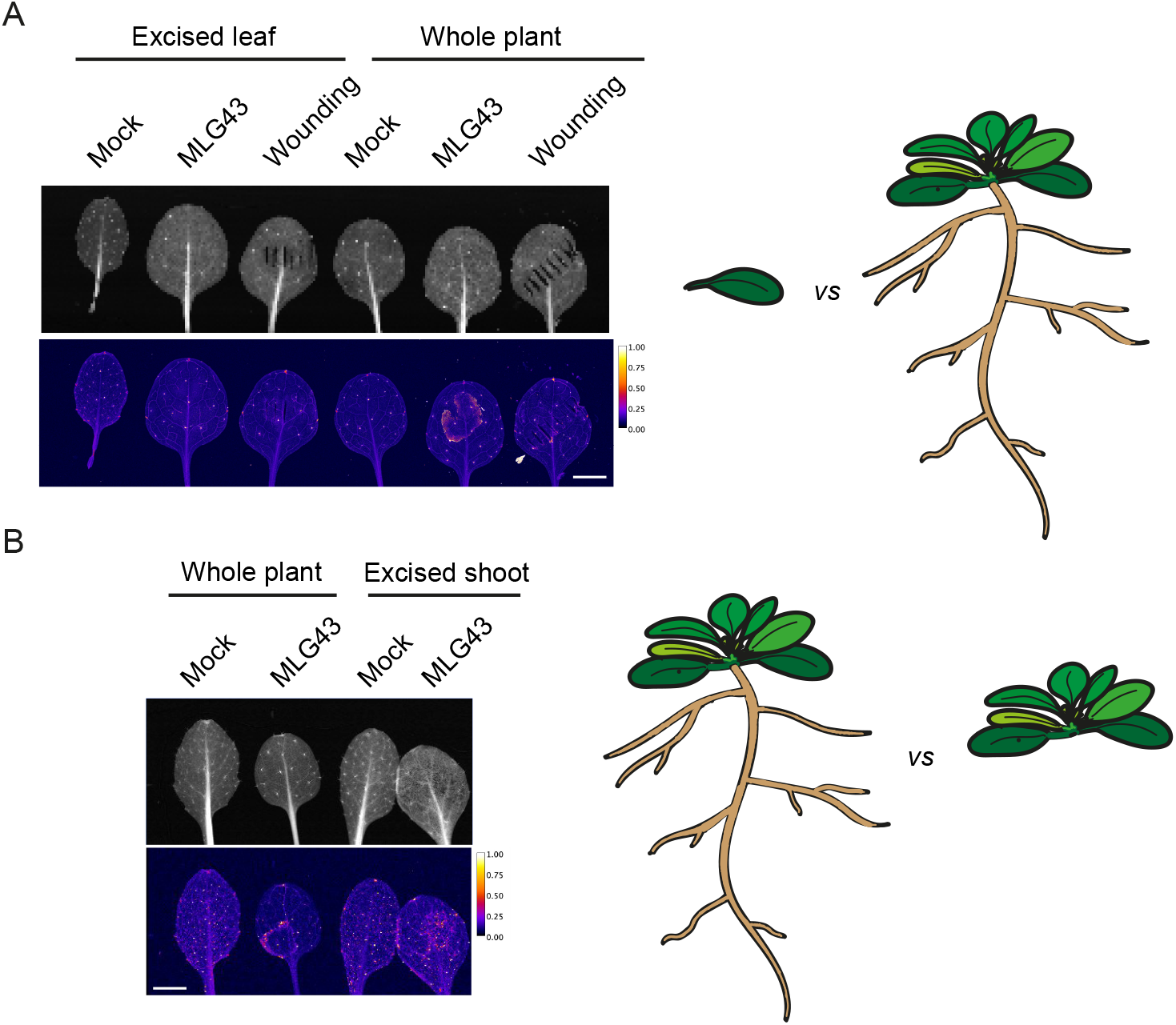
Apoplastic zinc accumulation requires distal resources. (A) Zinc accumulation in detached Col-0 leaves, excised before treatment and incubated for 24 h following application of a 5 µl-droplet of mock solution, 50 µM MLG43 or mechanical wounding with forceps. And zinc distribution in intact Col-0 rosettes, in which leaves remained attached and were exposed for 24 h a 5 µl-droplet of mock (water), 50 µM MLG43 or mechanical wounding. Upper panel shows transmission map. (B) S-XRF images of zinc distribution in treated leaves (mock and 50 µM MLG43) from plants with roots (whole plant) *versus* leaves from rosettes without roots (excised shoot). Units indicate areal density µg/cm^2^. Upper panel shows transmission map. Each image is representative of three independent replicates showing similar results. Scale bar =2 cm.

### 3.2. ZiMI configures a specific transcriptional program

To determine the molecular processes underlying ZiMI and its regulation, *Pc*BMM-induced zinc redistribution was determined in mutants impaired in the JA (*jar1-1*), ET (*ein2-5*), and SA (*sid2-1*) signalling pathways (Fig S1). In none of the observed mutants, zinc reallocation was fully abolished, indicating that pathogen-induced zinc accumulation does not strictly depend on JA-, SA-, or ET-mediated defence signalling.

Towards unveiling the genetic elements involved in ZiMI, RNA-Seq analyses were carried out in 16 days old wild type Col-0 and *hma2hma4* mutants mock-or *Pc*BMM-inoculated at 48hpi. Principal Component Analysis (PCA) mainly separated mock-treated Col-0 from the rest of the samples (PC1, 62% variance), being mock-treated *hma2hma4* grouped close to infected samples (Fig. 3A). Indeed, we identified 4,000-8,000 differentially expressed genes (DEGs; absolute log_2-_transformed fold-change ≥ 1, adjusted *P* ≤ 0.05, Wald’s test) in all pairwise comparisons with Col-0 under mock conditions (Fig. 3B). Quality control of the pairwise comparisons were done by observing large differences in the expression levels of genes involved in metal transport when comparing *hma2hma4* with Col-0 plants under mock conditions (3,790 DEGs; Table S1). As expected, metal transporters of the ZIP (Zinc-Regulated Transporter/ Iron-Regulated Transporter) family, such as *bZIP23* and *bZIP19,* typically involved in Zn^2+^ transport, together with nicotianamine synthase 2, *NAS2,* which contributes to long-distance delivery of Zn^2+^, were up-regulated by the loss of *HMA2* and *HMA4* (Lilay et al., 2021). According to Escudero et al., (2022) *hma2hma4* plants under mock conditions showed increased expression of the defence genes *PR1* and *LOX2* compared with mock-treated Col-0 plants. Interestingly, the *hma2hma4* mutants display constitutive activation of numerous defence-related genes what is consistent with its proximity to infected samples in the PCA analyses (Fig. 3A; Table S1). However, *hma2hma4* lacking these zinc transporters remained highly susceptible to *Pc*BMM (Escudero *et al*., 2022). Furthermore, and in line with previously observed in *Pc*BMM-inoculated plants, the fungal infection imposed upregulation of the transcript levels of *RbohD*, which encodes an NADPH oxidase involved in ROS production, along with defence-related genes such as *PR1, PDF1.2, PAD3*, and more than 40 members of the *WRKY* transcription factor family in Col-0 plants (4643 DEG; Table S2; Munoz-Barrios et al., 2020).

**Fig. 3.**
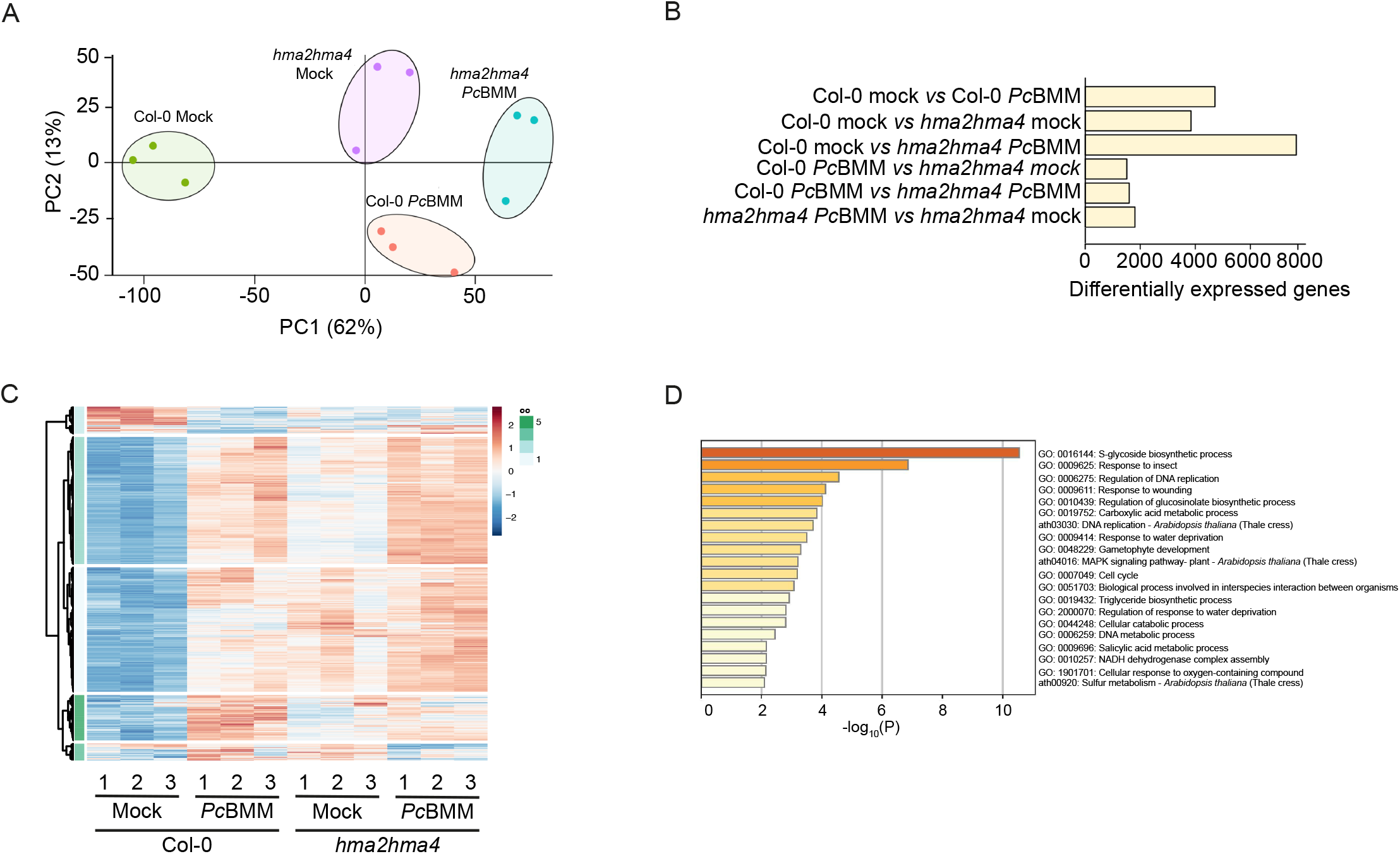
ZiMI induces a complex transcriptional reprograming. (A) Principal component analysis (PCA) of the three biological replicates for each genotype (Col-0 and *hma2hma4*) and each treatment (mock and *Pc*BMM inoculation) used for the RNA-sequencing. (B) Number of differentially expressed genes (DEGs) in all the possible comparisons. (C) Hierarchical clustering of 1467 DEGs due to the interaction of factors (genotype and infection). (D) Biological process gene ontology (GO) term enrichment bar graph of the 533 DEGs in the ZiMI regulon, coloured by *p*-values. Log10(P) is the *p*-value in log base 10 (Metascape).

To capture the most relevant transcriptome changes, z-scores of normalised counts of DEGs according to the interaction between genotype and treatment (adjusted *P* ≤ 0.05, Wald’s test, 1,467 DEG) were used to draw a hierarchical clustering heatmap according to the Ward D2 method (Fig. 3C). Five main trajectories could be envisaged (clusters I to V) (Fig. 3C; Table S3). Remarkably, functional analysis of transcripts within each cluster revealed significant enrichment in Gene Ontology (GO) terms for biological processes associated with plant immunity (adjusted *P* ≤ 0.05, Fisher’s Exact test), including defence responses to fungal and bacterial pathogens, PTI-triggered signalling, systemic acquired resistance, glucosinolate biosynthesis, and salicylic acid-and jasmonic acid-mediated responses among others. Overall, the lowest steady-state levels of the transcripts were found under mock-treated Col-0 and tended to raise upon infection, reflecting the triggering of the immune response. The trajectory of the transcripts included in clusters I and III (1,084 transcripts, 74% DEGs) reflected higher basal levels of immune-related features in mock-treated *hma2hma4* (more pronounced in cluster III compared to cluster I) that achieved an even more prominent accumulation upon the infection. This pattern is consistent with constitutive defence activation in the *hma2hma4* mutant. Conversely, *hma2hma4* failed to induce the transcripts in clusters IV and V (267 transcripts, 18% DEGs) to the levels observed in Col-0 following infection. This indicates an impaired activation of this defence-associated transcriptional programme, particularly genes involved in glucosinolate biosynthesis. In contrast, transcripts in cluster II (117 transcripts, 8% DEGs) were down-regulated in Col-0 following *Pc*BMM infection, but they remained at similar low basal expression levels in the *hma2hma4* mutant, showing little or no transcriptional response to infection. This set of DEGs was not significantly enriched for any Gene Ontology biological process. Thus, the transcriptome analysis supports a key role for HMA2 and HMA4 in the timely activation of immune responses against pathogens.

To identify the genes specifically involved in ZiMI, the DEGs upon *Pc*BMM infection (Col-0 *vs. hma2hma4* inoculated with *Pc*BMM; 1574 DEGs; Table S4) and those differing under mock conditions (Col-0 *vs. hma2hma4* mock; 3790 DEGs; Table S1) were compared to each other, selecting the 533 DEGs that only were differentially expressed in pathogenesis (Table S5). Gene ontology analyses of these genes indicated that they were largely associated with glucosinolate biosynthesis, 2-oxocarboxylic acid metabolism, response to insect and wounding and MAPK signalling pathway (including plasma membrane-associated receptor kinases) (Fig. 3D). This is consistent with a coordinated stress and defence-associated transcriptional reprogramming. Interestingly, only one metal (zinc) transporter gene, *ZIP2*, was in the ZiMI transcriptome, being among the 15 genes most highly expressed in Col-0 than *hma2hma4* (Table 1). At the other side of the spectrum, *NRAMP3* is the only metal transporter gene to be more expressed in infected *hma2hma4* plants than in Col-0 among the ZiMI genes. Only 9 genes within the ZiMI regulon indicated a change in the direction of the induction, moving from being more highly expressed in *hma2hma4* under mock conditions to being more expressed in Col-0 during infection (Table S5). However, no genes displayed the opposite pattern.

**Table 1.**
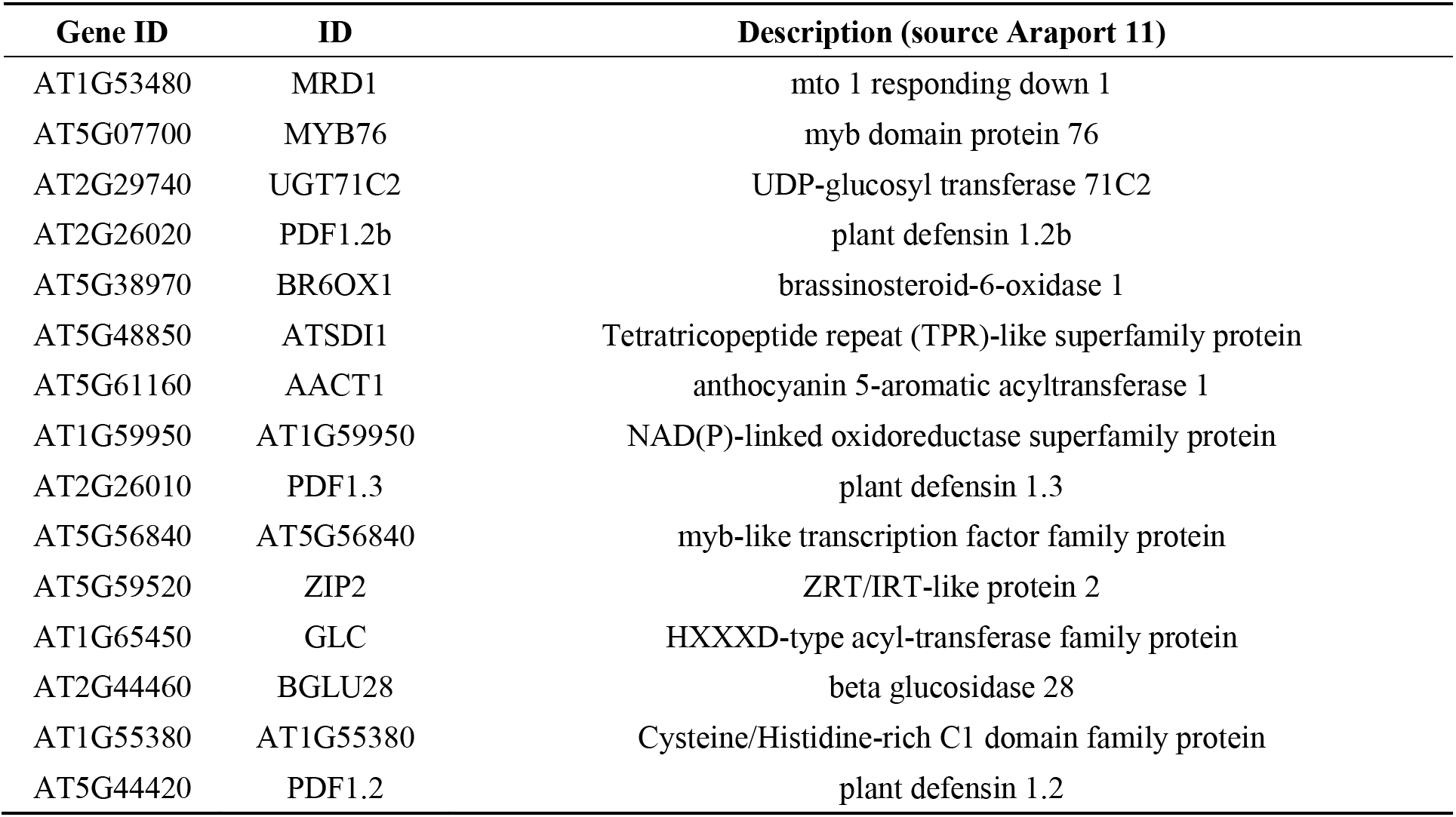
The 15 most highly expressed genes in the ZiMI transcriptome of Col-0 plants compared with the *hma2hma4* mutant.

Since ZiMI is strongly triggered by MLG43, the ZiMI DEGs were compared with those from a previous transcriptomic analysis in which Arabidopsis seedlings were treated for 30 min with 50 µM MLG43 (Rebaque *et al*., 2021). Despite being both studies conducted under different methodological approaches, 20 DEGs were shared between the two datasets (Table 2), with 17 of them maintaining the same direction of induction. As expected, these genes were associated with plant immunity, response to stress, and transport processes, although some of them are still uncharacterized genes. While no direct zinc homeostasis gene was observed, one of them (At2g44840, *ERF13*) has been shown to participate in zinc metabolism, as evidenced by the down-regulation of zinc transporter *ZIP11* and transcription factor *ZAT11* in *erf13* mutants (Chen et al., 2024).

**Table 2.**
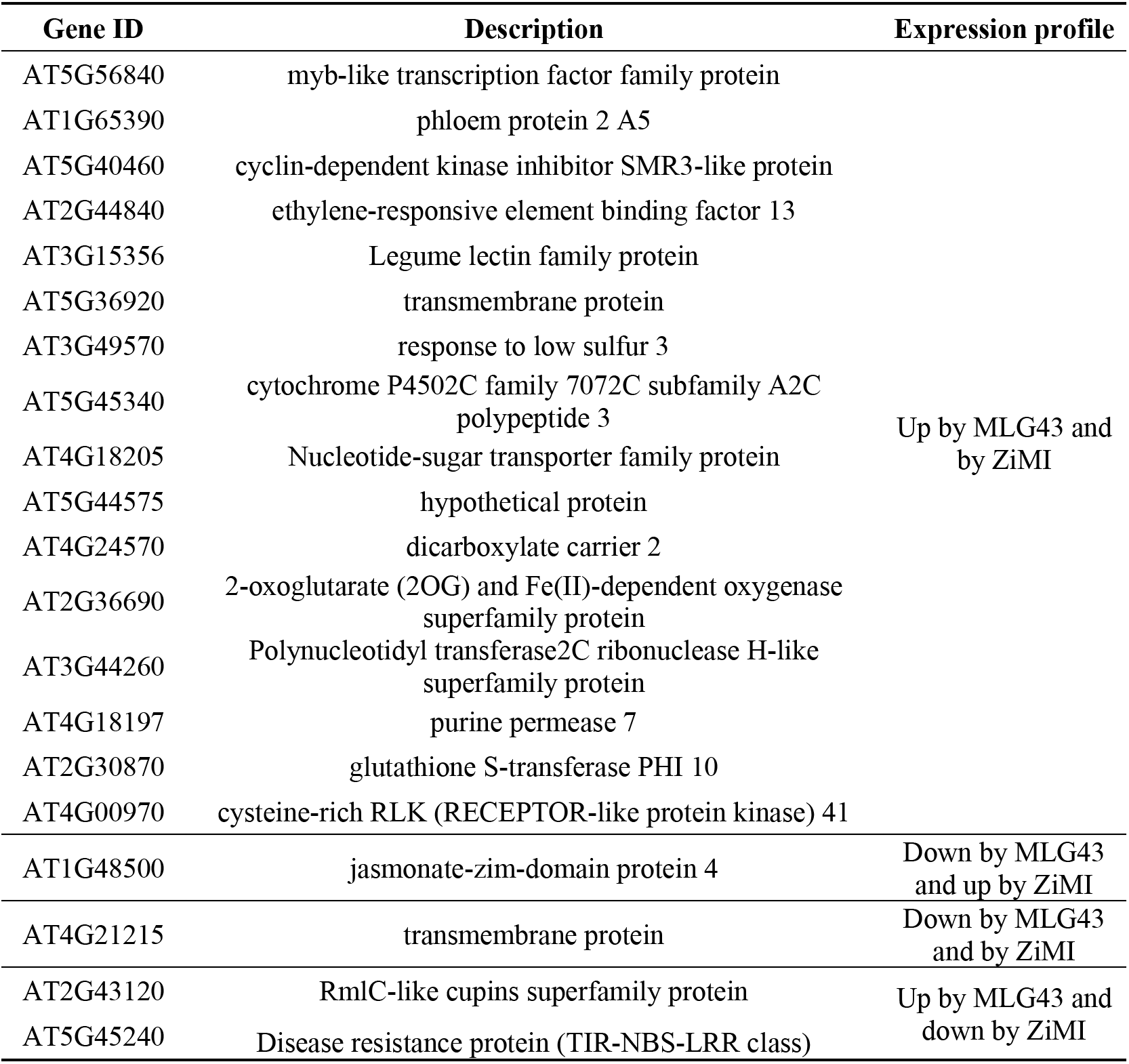
DEG shared by mock *vs* MLG43 6 h treatment and ZiMI. The table lists the 20 genes differentially expressed in both datasets. DEGs were defined as transcripts with |log2 fold-change| ≥ 1 and adjusted P ≤ 0.05 (Wald’s test).

### 3.3. Plant-synthesized metabolites in response to ZiMI

To determine the metabolic changes associated with ZiMI, un-targeted metabolomics approach was used with the same experimental conditions used for the transcriptional studies. In total, 466 different metabolites were detected (Table S6), the abundances of which were compared pairwise between plant lines and treatments (i.e., *Pc*BMM vs. mock) respectively (Fig 4A). The most pronounced differences in metabolite accumulation were observed between the plant genotypes of the same experimental condition. When comparing *Pc*BMM-inoculated to Col-0 plants, 9 metabolites were detected in *hma2hma4* exclusively, 31 at significantly higher and 20 at lower levels (Fig. 4B, C). For the mock-inoculated plants, 10 metabolites were found exclusively and 14 more abundantly in the zinc transport deficient *hma2hma4* lines, while another 12 metabolites were more abundant and 6 exclusively detected in mock-treated Col-0 plants. Less pronounced differences were found as a consequence of the pathogen inoculation (Fig. 4B, C).

**Fig. 4.**
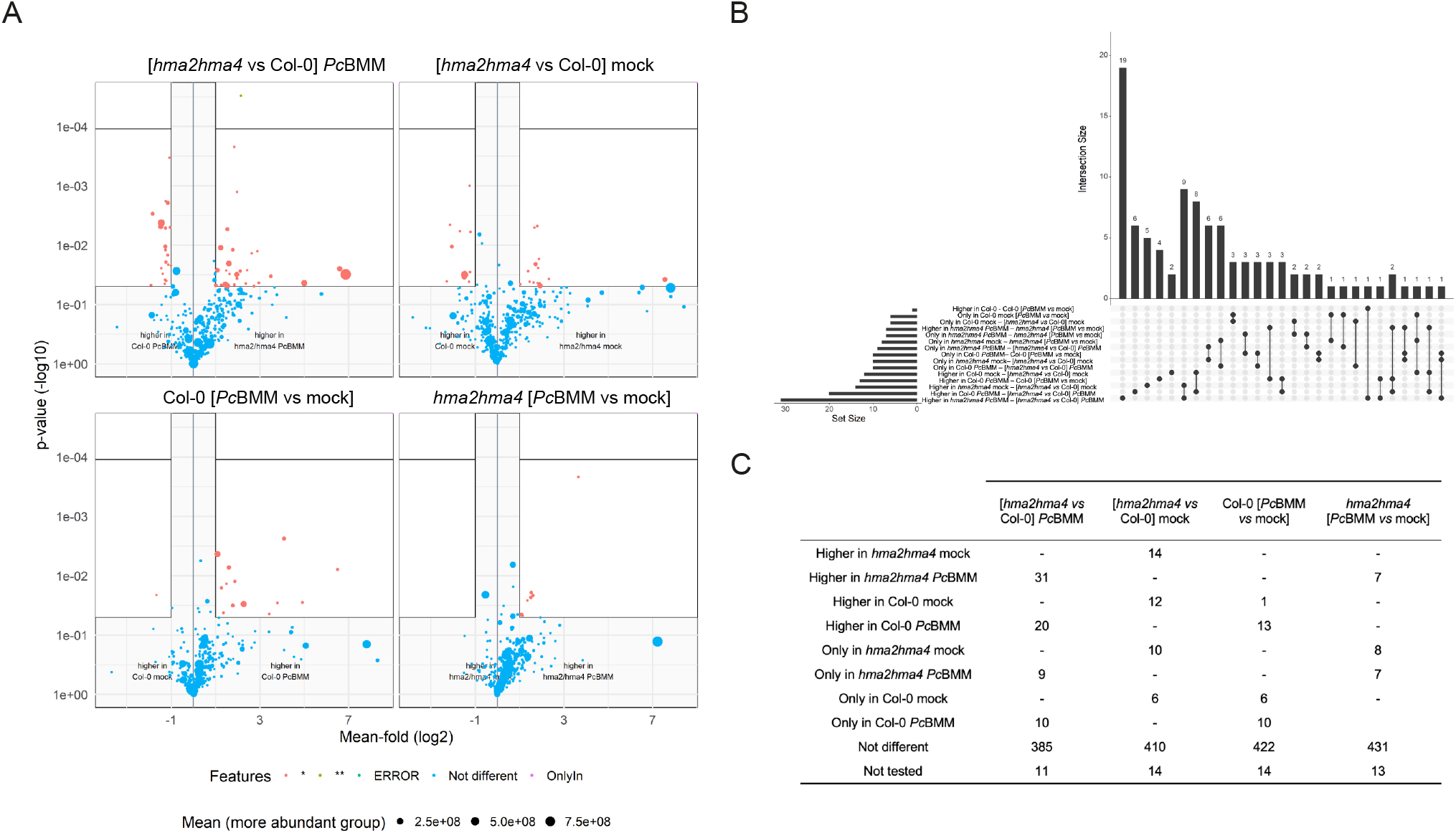
Mutations in *HMA2* and *HMA4* impact on specialized metabolites biosynthesis. (A) Volcano plots of the 4 univariate comparisons carried out for the dataset (B) The upset plot shows the aggregated results of the four univariate comparisons. The labels of the comparisons are in the form “<outcome> - <comparison>”, with outcome indicating the experimental group the metabolite was only detected in or more abundant in the respective comparison. The numbers of the “Set Size” and “Intersection Size” refer to metabolites being differently abundant in one or more comparisons. (C) Number of metabolites differentially produced in the pairwise comparisons.

By comparing the differences between the *Pc*BMM-inoculation and mock-treatment, four major clusters could be identified: those involved in a general immune process (significant changes only observed in infected *vs* mock, but not between genotypes; Table S7), including jasmonic acid and some indol-glucosinolates. Those participating in or affected by disturbed zinc homeostasis (the differences were only observed between genotypes, but not upon *Pc*BMM inoculation; Table S8), that involve nitrogen and phenylpropanoid related compounds that are detected at higher levels in the *hma2hma4* plants. Those linked to immunity, but with levels already higher in the mock-inoculated double mutant plants (Table S9); and those related to ZiMI (Table 3). The latter cluster involve those metabolites with altered synthesis during infection in Col-0 plants that at the same time present a significantly different abundance in infected *hma2hma4* plants and, consequently, would lead to an enhanced susceptibility against pathogens. Interestingly, among the 11 metabolites in this category (Table 3), the ones identified corresponds to feruloyl-agmatine and one isomer of coumaroyl-agmatine, known phenylpropanoid-derived phytoalexins involved in defence against pathogens and cell wall reinforcement (Keller et al., 1996; Liu et al., 2022; van Zadelhoff et al., 2022; Xue et al., 2025). Consistent with this, At5g61160 encoding the enzyme agmatine coumaroyl transferase AtACT required for the synthesis of this metabolite (Muroi et al., 2009), is down-regulated in *hma2hma4* and it is among the ZiMI regulon genes (Table S5). Notably, an octadecanoid compound (FP1627; Table 3) that is produced by lower levels during infection in Col-0 plants, is detected in higher amounts in infected *hma2hma4* double mutant than wild type plants. Similarly, *Pc*BMM-infected *hma2hma4* mutants produce significantly lower amounts of one metabolite represented by feature FP1508 that is associated with building physical barriers to prevent pathogen colonization (Table S6). FP1508 is associated with 3-hydroxy-palmitate, required for the synthesis of very long-chain fatty acids used for plant cuticle biosynthesis (Roudier et al., 2010). Therefore, transcriptomic and metabolomic data set underscore the multifaceted nature of ZiMI, associated with the alteration of phenylpropanoid metabolism, as well as the reinforcement of physical barriers through cell wall remodelling and cuticle reinforcement.

**Table 3.** ZiMI metabolites. Leve1 1: identified compound with authentic reference standard (RT, MZ, MSMS); Level 3: Putative substance classification (MS and MSMS); Level 5: Unequivocal molecular formula (MS isotope/adduct); Annotation levels according to (Schymanski et al., 2014)

| Num | O-Group | Putative sum formula | Significant Col-0 [PcBMM vs mock] | Significant PcBMM [*hma2hma4* vs Col-0] | Annotation type: Annotation/Identification |
| --- | --- | --- | --- | --- | --- |
| Fp.331 | Met.183 | C<sub>14</sub>H<sub>20</sub>N<sub>4</sub>O<sub>2</sub> | Higher in Col-0 *PcBMM* | Higher in Col-0 *PcBMM* | Level 1: Trans-Coumaroyl-Agmatine |
| Fp.372 | Met.190 | C<sub>15</sub>H<sub>22</sub>N<sub>4</sub>O<sub>3</sub> | Higher in Col-0 *PcBMM* | Higher in Col-0 *PcBMM* | Level 1: Trans-Feruloyl-Agmatine |
| Fp.1627 | Met.377 | C<sub>18</sub>H<sub>34</sub>O<sub>6</sub> | Higher in Col-0 mock | Higher in *hma2hma4* *PcBMM* | Level 3: Putative class Octadecanoid |
| Fp.108 | Met.355 | C<sub>11</sub>H<sub>14</sub>O | Only in Col-0 *PcBMM* | Only in Col-0 *PcBMM* | Level 5: sum formula from [M+H]<sup>+</sup> |
| Fp.1607 | Met.443 | C<sub>18</sub>H<sub>34</sub>O<sub>3</sub> | Only in Col-0 *PcBMM* | Only in Col-0 *PcBMM* | Level 5: sum formula from [M-H]<sup>-</sup> |
| Fp.29 | Met.170 | C<sub>10</sub>H<sub>8</sub>O<sub>3</sub> | Only in Col-0 *PcBMM* | Higher in *hma2hma4* *PcBMM* | Level 5: sum formula from [M+H]<sup>+</sup> |
| Fp.1034 | Met.61 | C<sub>7</sub>H<sub>16</sub>N<sub>4</sub>O | Higher in Col-0 *PcBMM* | Higher in Col-0 *PcBMM* | Level 5: sum formula from [M+H]<sup>+</sup> |
| Fp.787 | Met.436 | C<sub>37</sub>H<sub>56</sub>O<sub>7</sub> | Higher in Col-0 *PcBMM* | Higher in Col-0 *PcBMM* | Level 5: sum formula from [M+Na]<sup>+</sup> |
| Fp.1503 | Met.416 | C<sub>16</sub>H<sub>28</sub>O<sub>3</sub> | Only in Col-0 mock | Only in *hma2hma4* *PcBMM* | Level 5: sum formula from [M-H]<sup>-</sup> |
| Fp.1220 | Met.23 |  | Higher in Col-0 *PcBMM* | Higher in Col-0 *PcBMM* |  |
| F.1516 | Met.467 |  | Only in Col-0 *PcBMM* | Only in Col-0 *PcBMM* |  |

## 4. Discussion

Zinc is an essential nutrient typically used as a cofactor in a large number of proteins. More recently, it has become apparent that zinc can act as a signal to induce developmental programs in plants, and a key element of plant immune response (Lin et al., 2024; Stanton et al., 2022). Zinc accumulation at infection sites has been reported in a number of species (De et al., 2025). ZiMI is a defence mechanism within PTI triggered upon MAMP/DAMPs perception. Here we demonstrated that Arabidopsis plants accumulated zinc in response to the β-1,3/1,4-mixed-linked glucan-derived oligosaccharide MLG43, that acts as a MAMP in this plant species (Rebaque et al., 2021). Zinc concentration at the inoculation site was not the result of cellular damage; cell integrity was maintained after MLG43 application, as indicated by potassium being confined within the cell. Furthermore, the zinc used in ZiMI seemed not to originate from the cells in the inoculation area, but from other leaves in the rosette. Only when the rosette was intact, zinc levels increased at the infection site. This is reminiscent on how nutrients are recovered from infected leaves, in which YSL1 and YSL3 transporters are upregulated to mobilize iron, copper, and zinc towards sink tissues such as newer leaves or flowers (Waters et al., 2006). In fact, infection with *Pc*BMM also induced *YSL3* transcription, what could reflect how zinc gets recovered from the neighbour leaves. Together, these findings suggest that zinc mobilization from adjacent tissues requires a coordinated response that is precisely triggered upon MAMP perception. While ZiMI was mostly observed with MLG43 application, we cannot exclude the involvement of other DAMPs or MAMPs in ZiMI activation as zinc-imaging was performed at a single time point, and the kinetics of signal transduction and of zinc efflux are likely to differ among elicitors.

The emerging relevance of MLG-derived oligosaccharides in plant-microbe interactions is underscored by the identification of multiple receptors involved in their perception across different plant species. In rice, MLG perception has been associated with the LysM receptor complex OsCERK1–OsCEBiP and the lectin receptor kinase OsLecRK1 (Dai et al., 2023; Yang et al., 2021). In Arabidopsis, the LysM receptor kinases CERK1, LYK4, and LYK5, and more recently the LRR-malectin receptor kinases IGP1 and IGP4, may contribute to MLG43 perception (Fernández-Calvo et al., 2024; Rebaque et al., 2021). Our results indicate that zinc accumulation upon MLG43 perception was partially dependent on the LysM receptor kinases, as *cerk1-2ly4lyk5* plants exhibited a reduced zinc distribution in the treated leaf area, but the response was not fully abolished. While the triple mutant showed a partial impairment, the *igp1* and *igp4* mutants showed no significant alteration. This difference suggests that some functional redundancy between *IGP* family could mask their role, a possibility that warrants further experimental testing.

MLGs present in some plant cell walls and microbial surfaces trigger rapid immune responses such as cytoplasmic Ca^2+^ fluxes, reactive oxygen species production, activation of mitogen-activated protein kinases cascades, leading to disease resistance in plants (Rebaque et al., 2025; Rebaque et al., 2021). Consistent with its elicitor role, MLG43 treatment up-regulated the expression of the zinc transporters *HMA2* and *HMA4*. However, this response appears to be independent of SA, JA and ET-mediated signalling, as *jar1-1*, *sid2-1* and *ein2-5* mutants retained the ability to actively export zinc to the apoplast upon pathogen challenge. These results suggest the existence of a ZiMI transcriptional program independent of the classical hormone-dependent ones.

Comparative transcriptomic analysis among mock-or *Pc*BMM-inoculated Col-0 or *hma2hma4* plants revealed large transcriptional reprogramming. One striking observation was the constitutive up-regulation of a large number of defence-related genes in the *hma2hma4* plants already detected in the mock-control. These included WRKY transcription factors (39 genes), cystein-rich receptor-like kinases (CRKs, 24 genes), and disease resistance and LRR proteins. Theoretically, this should lead to *hma2hma4* plants being more resistant to disease. However, the mutant was more susceptible to the necrotrophic fungus *Pc*BMM (Escudero et al., 2022). The precise cause for these contrasting observations must lie on the altered zinc status of *hma2hma4* plants. Lack of zinc would trigger the upregulation of genes involved in the immune response through three different mechanisms: i) zinc is used a cofactor of some of these enzymes, and the organism reacts to their loss of activity by inducing their transcription, ii) absence of zinc can be sensed as a pathogen removing it, triggering an immune response, or iii) a zinc-dependent protein is acting as repressor of the expression of some of these genes. In any case, the increased susceptibility of *hma2hma4* mutants highlights the importance of zinc as an essential element in plant-immunity. Considering the low zinc bioavailability in large areas of the world (Alloway, 2008), zinc fertilization should not be only targeted to seed biofortification, but also to ensure optimized plant immune responses.

The RNA-seq analysis revealed an extensive transcriptional reprograming and allowed the identification of a ZiMI regulon. Overall, ZiMI involved the expression of genes associated with activation of toxic mechanisms against pathogens, danger perception and signalling, and reinforcement of apoplastic barriers. These included signalling components (16 kinases and 7 receptors), lipid transfer proteins and genes involved in secondary metabolism and cell wall remodelling, and defensins. Beyond its well-established antimicrobial activity against a broad range of microorganisms (Thomma et al., 2002), the coordinated induction of multiple defensins may also be relevant in the context of zinc homeostasis (Van der Weerden & Anderson 2013). Their high cysteine content raises the possibility that some members may contribute to local zinc binding or buffering within the apoplast, thereby modulating zinc availability during immune responses. Interestingly, some of the the ZiMI-induced genes included receptor proteins such as cysteine-rich receptor like kinase 41 (CRK41) that contained a putative Zn^2+^-binding motif. It is tempting to speculate with a putative role of apoplastic Zn^2+^ in triggering CRK41. This deserves further investigation since it may represent a mechanism linking zinc homeostasis to the propagation of plant immune signalling.

The increased susceptibility of *hma2hma4* to *Pc*BMM must emanate from a few metabolites and enzymes that are not produced at sufficient levels during infection due to altered zinc allocation. Our untargeted metabolomics approach suggests that molecules involved in plant defence against pathogens may contribute to ZiMI. Among them are hydroxycinnamic acid amides (HCAAs), isoforms of feruoyl-agmatine and coumaroyl-agmatine. These metabolites are widely distributed across plant species, and they have direct antimicrobial activity (Carere et al., 2018; Muroi et al., 2009). Furthermore, they play critical roles in plant physiology, including structural cell wall reinforcement in abiotic/biotic stress responses (Xue et al., 2025). Notably, the *AtACT* gene, that mediates the synthesis of these two molecules belongs to the ZiMI regulon. Moreover, it is not only upregulated in response to *Pc*BMM infection, but also to other necrotrophic fungi such as *Alternaria brassicicola* infection, supporting its relevant role in defence (Munoz-Barrios et al., 2020; Muroi et al., 2009).

A closer look at the differentially accumulated metabolites and their putative annotations (level 3, no identification) pointed to a central role for phenylpropanoid metabolism in ZiMI. Beyond the antimicrobial activity, the phenylpropanoid pathway also contributes to plant defence by reinforcing the cell wall through the deposition of macromolecular phenolic structures by covalent linkage of compounds such as the HCAAs and suberin. Additionally, 3-hydroxypalmitic acid can contribute to the strength of physical barriers against pathogen invasion, as an intermediate involved in the biosynthesis of cuticular waxes and suberin. Together, these metabolites define a coherent metabolic signature of defence activation. Consistent with these metabolic changes, the transcriptomic analysis revealed the induction of key genes involved in phenylpropanoid biosynthesis, including phenyl ammonia lyase 1(*PAL1*) and coumarate::CoA ligase (*4CL*), as well as genes associated with lipid metabolism and cuticle reinforcement, such a long-chain fatty acid alcohol dehydrogenase, lipid transfer proteins and *CYP77A6*. Considering that their absence is associated with *hma2hma4* being more susceptible despite the overproduction of other molecules and specialized metabolites involved in plant immunity, we can conclude that these metabolites play a critical, central role in Zn-mediated immunity. Notably, the ZiMI regulon also contains a core of genes involved in aliphatic glucosinolate biosynthesis, such as *CYP79F1*, *CYP79F2* and *CYP83A1*, suggesting the coordinated activation of multiple defence layers during this immune response.

## 5. Conclusion

In this study we have combined elemental imaging, transcriptomic and metabolomic analyses to functionally dissect a previously uncharacterized layer of plant immunity named ZiMI. Locally increasing zinc to toxic levels is part of a coordinated PTI response. ZiMI does not only act directly on the pathogen but it also triggers a coordinated reprogramming of glucosinolate and phenylpropanoid metabolism, increasing the flow towards chemical and structural strengthening of the cell wall. These findings expand the knowledge of PTI mechanisms and provide new molecular tools for improving resistance against pathogens.

## Supporting information

Additional supporting information may be found online in the Supporting Information section at the end of the article.

## CRediT authorship contribution statement

**Viviana Escudero:** investigation, data curation, writing-review and editing. **Cuong V. Hoang**: investigation. **Antoni García-Molina:** data curation, formal analysis, writing-review and editing. **Aishee De**: investigation. **Alejandro M. Armas**: investigation. **Dennis Brueckner**: investigation, data curation, formal analysis, writing-review and editing. **DFS**: investigation, data curation, formal analysis, writing-review and editing. **Christoph Bueschl**: data curation, formal analysis, writing-review and editing. **Maria Doppler**: data curation, formal analysis. **Antony van de Ent:** data curation, formal analysis. **Rainer Schuhmacher:** data curation, formal analysis, writing-review and editing. **Manuel González-Guerrero:** conceptualization, writing original draft, supervision, data curation, review and editing, funding acquisition. **Lucía Jordá:** conceptualization, writing original draft, supervision, data curation, review and editing, funding acquisition

## Acknowledgments

This work was financially supported by grant TED2021-131769B-I00 to LJ/MGG funded by MICIU/AEI/10.13039/501100011033 and by “European Union NextGenerationEU//PRTR, and Severo Ochoa Program for Centres of Excellence in R&D (grant SEV-2016-0672 and CEX2020-000999-S (2022-2025)) funded by MICIU/AEI/10.13039/501100011033. VE, CVH, and AD were recipient of postdoctoral contracts funded by MCIN/AEI/10.13039/501100011033. AG-M is recipient of the grant RYC2022-037020-I funded by MICIU/AEI/ 10.13039/501100011033 and by “ESF+”. We also acknowledge financial support from the MICIU/AEI/10.13039/501100011033 through the “Severo Ochoa Program for Centres of Excellence in R&D” (CEX2019-000902-S) and the CERCA Program/Generalitat de Catalunya. AMA is funded by MSCA grant HORIZON-MSCA-2022-PF-01-101104098-ZINMUNITY. We acknowledge DESY (Hamburg, Germany), a member of the Helmholtz Association HGF, for the provision of experimental facilities. Parts of this research were carried out at PETRA III. Data were collected using beamline P06 operated/provided by DESY Photon Science. Beamtime was allocated for proposals I-20230698 EC, and I-20250029. This research was supported in part through the Maxwell computational resources operated at Deutsches Elektronen-Synchrotron DESY, Hamburg, Germany. We acknowledge the Paul Scherrer Institute (Villigen, Switzerland) for beamtime in line microXAS through proposal 20221807. The HRMS equipment for metabolomics measurements was kindly provided by the BOKU Core Facility Bioactive Molecules: Screening and Analysis (RRID:SCR_028285).

## Supplementary materials

Supplementary material associated with this article can be found, in the online version.

**Table S1. DEGs in *hma2hma4* under mock conditions**

**Table S2. DEGs in Col-0 upon *Pc*BMM infection**

**Table S3. DEGs associated with the genotype x treatment interaction**

**Table S4. DEGS in *hma2hma4* upon *Pc*BMM infection**

**Table S5. ZiMI regulon**

**Table S6. Differential features detected by untargeted metabolomics**

**Table S7. Metabolites affected by infection and not genotype**

**Table S8. Metabolites affected by mutation and nor infection**

**Table S9. Preactivated immune metabolites in *hma2hma4***

**Table S10. Primers used**

## Data availability

RNAseq were deposited at the Gene Expression Omnibus (GEO).

---

**Fig. S1. PTI-triggered zinc accumulation is independent of the JA, SA or ET-mediated signalling pathways.** Synchrotron-based X-ray fluorescence images of wild type Col-0, *jar1-1*, *sid2-1* and *ein2-5* mutants at 48 hpi under mock or *Pc*BMM-spore inoculation conditions. Left-most column shows the calcium distribution upon mock treatment, next to it is zinc distribution upon mock; followed by Ca and Zn localization after *Pc*BMM droplet inoculation. Units indicate areal density µg/cm^2^. Scale bars = 1 mm. Images were taken from randomly chosen leaves. At least five independent plants from the indicated genotypes were used for each treatment.

## Notes

### Competing Interest Statement

The authors have declared no competing interest.

## Reference

Alloway, B.J. (2008) Zinc in Soils and Crop Nutrition, 2nd ed. published by IZA and IFA, Brussels, Belgium and Paris, France

Bolger, A.M., Lohse, M., and Usadel, B. (2014). Trimmomatic: a flexible trimmer for Illumina sequence data. Bioinformatics 30:2114–2120. 10.1093/bioinformatics/btu170.

Bueschl, C., Kluger, B., Neumann, N.K.N., Doppler, M., Maschietto, V., Thallinger, G.G., Meng-Reiterer, J., Krska, R., and Schuhmacher, R. (2017). MetExtract II: A Software Suite for Stable Isotope-Assisted Untargeted Metabolomics. Anal Chem 89:9518–9526. 10.1021/acs.analchem.7b02518.

Cabot, C., Martos, S., Llugany, M., Gallego, B., Tolrà, R., and Poschenrieder, C. (2019). A Role for Zinc in Plant Defense Against Pathogens and Herbivores. Front Plant Sci 10:1171. 10.3389/fpls.2019.01171.

Campos, M.L., de Souza, C.M., de Oliveira, K.B.S., Dias, S.C., and Franco, O.L. (2018). The role of antimicrobial peptides in plant immunity. J Exp Bot 69:4997–5011. 10.1093/jxb/ery294.

Cao, M., Platre, M.P., Tsai, H.H., Zhang, L., Nobori, T., Armengot, L., Chen, Y., He, W., Brent, L., Coll, N.S., et al. (2024). Spatial IMA1 regulation restricts root iron acquisition on MAMP perception. Nature 625:750–759. 10.1038/s41586-023-06891-y.

Chen, W., Shi, Y., Wang, C., and Qi, X. (2024). AtERF13 and AtERF6 double knockout fine-tunes growth and the transcriptome to promote cadmium tolerance in Arabidopsis. Gene 911:148348. 10.1016/j.gene.2024.148348.

Chinchilla, D., Boller, T., and Robatzek, S. (2007). Flagellin signalling in plant immunity. Adv Exp Med Biol 598:358–371. 10.1007/978-0-387-71767-8_25.

Couto, D., and Zipfel, C. (2016). Regulation of pattern recognition receptor signalling in plants. Nat Rev Immunol 16:537–552. 10.1038/nri.2016.77.

Dai, Y.S., Liu, D., Guo, W., Liu, Z.X., Zhang, X., Shi, L.L., Zhou, D.M., Wang, L.N., Kang, K., Wang, F.Z., et al. (2023). Poaceae-specific β-1,3;1,4-d-glucans link jasmonate signalling to OsLecRK1-mediated defence response during rice-brown planthopper interactions. Plant Biotechnol J 21:1286–1300. 10.1111/pbi.14038.

Dangol, S., Chen, Y., Hwang, B.K., and Jwa, N.S. (2019). Iron-and Reactive Oxygen Species-Dependent Ferroptotic Cell Death in Rice-Magnaporthe oryzae Interactions. Plant Cell 31:189–209. 10.1105/tpc.18.00535.

De, A., Hoang, C.V., Escudero, V., Armas, A.M., Echavarri-Erasun, C., Gonzalez-Guerrero, M., and Jorda, L. (2024). Combating plant diseases through transition metal allocation. New Phytol 10.1111/nph.20366.

DeFalco, T.A., and Zipfel, C. (2021). Molecular mechanisms of early plant pattern-triggered immune signaling. Mol Cell 81:4346. 10.1016/j.molcel.2021.09.028.

Dobin, A., Davis, C.A., Schlesinger, F., Drenkow, J., Zaleski, C., Jha, S., Batut, P., Chaisson, M., and Gingeras, T.R. (2013). STAR: ultrafast universal RNA-seq aligner. Bioinformatics 29:15–21. 10.1093/bioinformatics/bts635.

Doppler, M., Bueschl, C., Ertl, F., Woischitzschlaeger, J., Parich, A., and Schuhmacher, R. (2022). Towards a broader view of the metabolome: untargeted profiling of soluble and bound polyphenols in plants. Anal Bioanal Chem 414:7421–7433. 10.1007/s00216-022-04134-z.

Duhrkop, K., Fleischauer, M., Ludwig, M., Aksenov, A.A., Melnik, A.V., Meusel, M., Dorrestein, P.C., Rousu, J., and Bocker, S. (2019). SIRIUS 4: a rapid tool for turning tandem mass spectra into metabolite structure information. Nat Methods 16:299–302. 10.1038/s41592-019-0344-8.

Escudero, V., Ferreira Sánchez, D., Abreu, I., Sopeña-Torres, S., Makarovsky-Saavedra, N., Bernal, M., Krämer, U., Grolimund, D., González-Guerrero, M., and Jordá, L. (2022). Arabidopsis thaliana Zn2+-efflux ATPases HMA2 and HMA4 are required for resistance to the necrotrophic fungus Plectosphaerella cucumerina BMM. J Exp Bot 73:339–350. 10.1093/jxb/erab400.

Fernández-Calvo, P., López, G., Martín-Dacal, M., Aitouguinane, M., Carrasco-López, C., González-Bodí, S., Bacete, L., Mélida, H., Sánchez-Vallet, A., and Molina, A. (2024). Leucine rich repeat-malectin receptor kinases IGP1/CORK1, IGP3 and IGP4 are required for arabidopsis immune responses triggered by β-1,4-D-Xylo-oligosaccharides from plant cell walls. Cell Surf 11:100124. 10.1016/j.tcsw.2024.100124.

Fuertes-Rabanal, M., Largo-Gosens, A., Fischer, A., Munzert, K.S., Carrasco-Lopez, C., Sanchez-Vallet, A., Engelsdorf, T., and Melida, H. (2024). Linear beta-1,2-glucans trigger immune hallmarks and enhance disease resistance in plants. J Exp Bot 75:7337–7350. 10.1093/jxb/erae368.

Goldstein, S., Meyerstein, D., and Czapski, G. (1993). The Fenton reagents. Free Radic Biol Med 15:435–445. 10.1016/0891-5849(93)90043-t.

Carere, J., Powell, J., Fitzgerald, T., Kazan, K., Gardiner, D.M. (2018). BdACT2a encodes an agmatine coumaroyl transferase required for pathogen defence in *Brachypodium distachyon*, Physiol Mol Plant Pathol. 104: 69–76. 10.1016/j.pmpp.2018.09.003.

Keller, H., Hohlfeld, H., Wray, V., Hahlbrock, K., Scheel, D., and Strack, D. (1996). Changes in the accumulation of soluble and cell wall-bound phenolics in elicitor-treated cell suspension cultures and fungus-infected leaves of *Solanum tuberosum*. Phytochemistry. 42:389–396.

Kind, T., and Fiehn, O. (2007). Seven Golden Rules for heuristic filtering of molecular formulas obtained by accurate mass spectrometry. BMC Bioinformatics 8:105. 10.1186/1471-2105-8-105.

Kramer, U. (2024). Metal Homeostasis in Land Plants: A Perpetual Balancing Act Beyond the Fulfilment of Metalloproteome Cofactor Demands. Annu Rev Plant Biol 75:27–65. 10.1146/annurev-arplant-070623-105324.

Kunze, G., Zipfel, C., Robatzek, S., Niehaus, K., Boller, T., and Felix, G. (2004). The N terminus of bacterial elongation factor Tu elicits innate immunity in Arabidopsis plants. Plant Cell 16:3496–3507. 10.1105/tpc.104.026765.

Kuvelja, A., Morina, F., Mijovilovich, A., Bokhari, S.N.H., Konik, P., Koloniuk, I., and Kupper, H. (2024). Zinc priming enhances Capsicum annuum immunity against infection by Botrytis cinerea-From the whole plant to the molecular level. Plant Sci 343:112060. 10.1016/j.plantsci.2024.112060.

Liao, Y., Smyth, G.K., and Shi, W. (2014). featureCounts: an efficient general purpose program for assigning sequence reads to genomic features. Bioinformatics 30:923–930. 10.1093/bioinformatics/btt656.

Lilay, G.H., Persson, D.P., Castro, P.H., Liao, F., Alexander, R.D., Aarts, M.G.M., and Assunção, A.G.L. (2021). Arabidopsis bZIP19 and bZIP23 act as zinc sensors to control plant zinc status. Nat Plants 7:137–143. 10.1038/s41477-021-00856-7.

Lin, J., Bjork, P.K., Kolte, M.V., Poulsen, E., Dedic, E., Drace, T., Andersen, S.U., Nadzieja, M., Liu, H., Castillo-Michel, H., et al. (2024). Zinc mediates control of nitrogen fixation via transcription factor filamentation. Nature 631:164–169. 10.1038/s41586-024-07607-6.

Liu, B., Li, J.F., Ao, Y., Qu, J., Li, Z., Su, J., Zhang, Y., Liu, J., Feng, D., Qi, K., et al. (2012). Lysin motif-containing proteins LYP4 and LYP6 play dual roles in peptidoglycan and chitin perception in rice innate immunity. Plant Cell 24:3406–3419. 10.1105/tpc.112.102475.

Liu, J., Li, W., Wu, G., and Ali, K. (2024). An update on evolutionary, structural, and functional studies of receptor-like kinases in plants. Front Plant Sci 15:1305599. 10.3389/fpls.2024.1305599.

Liu, S., Jiang, J., Ma, Z., Xiao, M., Yang, L., Tian, B., Yu, Y., Bi, C., Fang, A., and Yang, Y. (2022). The Role of Hydroxycinnamic Acid Amide Pathway in Plant Immunity. Front Plant Sci 13:922119. 10.3389/fpls.2022.922119.

Love, M.I., Huber, W., and Anders, S. (2014). Moderated estimation of fold change and dispersion for RNA-seq data with DESeq2. Genome Biol 15:550. 10.1186/s13059-014-0550-8.

Macomber, L., and Imlay, J.A. (2009). The iron-sulfur clusters of dehydratases are primary intracellular targets of copper toxicity. Proc Natl Acad Sci U S A 106:8344–8349. 10.1073/pnas.0812808106.

Martín-Dacal, M., Fernández-Calvo, P., Jiménez-Sandoval, P., López, G., Garrido-Arandía, M., Rebaque, D., Del Hierro, I., Berlanga, D.J., Torres, M., Kumar, V., et al. (2023). Arabidopsis immune responses triggered by cellulose-and mixed-linked glucan-derived oligosaccharides require a group of leucine-rich repeat malectin receptor kinases. Plant J 113:833–850. 10.1111/tpj.16088.

Molina, A., Sanchez-Vallet, A., Jorda, L., Carrasco-Lopez, C., Rodriguez-Herva, J.J., and Lopez-Solanilla, E. (2024a). Plant cell walls: source of carbohydrate-based signals in plant-pathogen interactions. Curr Opin Plant Biol 82:102630. 10.1016/j.pbi.2024.102630.

Molina, A., Jorda, L., Torres, M.A., Martin-Dacal, M., Berlanga, D.J., Fernandez-Calvo, P., Gomez-Rubio, E., and Martin-Santamaria, S. (2024b). Plant cell wall-mediated disease resistance: Current understanding and future perspectives. Mol Plant 17:699–724. 10.1016/j.molp.2024.04.003.

Morina, F., and Küpper, H. (2022). Trace metals at the frontline of pathogen defence responses in non-hyperaccumulating plants. J Exp Bot 73:6516–6524. 10.1093/jxb/erac316.

Morina, F., Mijovilovich, A., Koloniuk, I., Pěnčík, A., Grúz, J., Novák, O., and Küpper, H. (2021). Interactions between zinc and Phomopsis longicolla infection in roots of Glycine max. J Exp Bot 72:3320–3336. 10.1093/jxb/erab052.

Munoz-Barrios, A., Sopena-Torres, S., Ramos, B., Lopez, G., Del Hierro, I., Diaz-Gonzalez, S., Gonzalez-Melendi, P., Melida, H., Fernandez-Calleja, V., Mixao, V., et al. (2020). Differential Expression of Fungal Genes Determines the Lifestyle of Plectosphaerella Strains During Arabidopsis thaliana Colonization. Mol Plant Microbe Interact 33:1299–1314. 10.1094/MPMI-03-20-0057-R.

Murdoch, C.C., and Skaar, E.P. (2022). Nutritional immunity: the battle for nutrient metals at the host-pathogen interface. Nat Rev Microbiol 20:657–670. 10.1038/s41579-022-00745-6.

Muroi, A., Ishihara, A., Tanaka, C., Ishizuka, A., Takabayashi, J., Miyoshi, H., and Nishioka, T. (2009). Accumulation of hydroxycinnamic acid amides induced by pathogen infection and identification of agmatine coumaroyltransferase in Arabidopsis thaliana. Planta 230:517–527. 10.1007/s00425-009-0960-0.

Rebaque, D., Del Hierro, I., López, G., Bacete, L., Vilaplana, F., Dallabernardina, P., Pfrengle, F., Jordá, L., Sánchez-Vallet, A., Pérez, R., et al. (2021). Cell wall-derived mixed-linked β-1,3/1,4-glucans trigger immune responses and disease resistance in plants. Plant J 106:601–615. 10.1111/tpj.15185.

Rebaque, D., Carrasco-Lopez, C., Krishnan, P., Lopez, G., Lopez-Cobos, S., de Salas, F., Meile, L., Lorrain, C., Largo-Gosens, A., McDonald, B.A., et al. (2025). Zymoseptoria tritici stealth infection is facilitated by stage-specific downregulation of a beta-glucanase. New Phytol 248:3191–3207. 10.1111/nph.70621.

Roudier, F., Gissot, L., Beaudoin, F., Haslam, R., Michaelson, L., Marion, J., Molino, D., Lima, A., Bach, L., Morin, H., et al. (2010). Very-long-chain fatty acids are involved in polar auxin transport and developmental patterning in Arabidopsis. Plant Cell 22:364–375. 10.1105/tpc.109.071209.

Schmid, R., Heuckeroth, S., Korf, A., Smirnov, A., Myers, O., Dyrlund, T.S., Bushuiev, R., Murray, K.J., Hoffmann, N., Lu, M., et al. (2023). Integrative analysis of multimodal mass spectrometry data in MZmine 3. Nat Biotechnol 41:447–449. 10.1038/s41587-023-01690-2.

Schymanski, E.L., Jeon, J., Gulde, R., Fenner, K., Ruff, M., Singer, H.P., and Hollender, J. (2014). Identifying small molecules via high resolution mass spectrometry: communicating confidence. Environ Sci Technol 48:2097–2098. 10.1021/es5002105.

Singh, G., Agrawal, H., and Bednarek, P. (2023). Specialized metabolites as versatile tools in shaping plant-microbe associations. Mol Plant 16:122–144. 10.1016/j.molp.2022.12.006.

Snoeck, S., Johanndrees, O., Nurnberger, T., and Zipfel, C. (2025). Plant pattern recognition receptors: from evolutionary insight to engineering. Nat Rev Genet 26:268–278. 10.1038/s41576-024-00793-z.

Stanton, C., Sanders, D., Kramer, U., and Podar, D. (2022). Zinc in plants: Integrating homeostasis and biofortification. Mol Plant 15:65–85. 10.1016/j.molp.2021.12.008.

Sun, G., Xiao, Y., Yin, H., Yu, K., Wang, Y., and Wang, Y. (2025). Oligosaccharide elicitors in plant immunity: Molecular mechanisms and disease resistance strategies. Plant Commun:101469. 10.1016/j.xplc.2025.101469.

Thomma, B.P., Cammue, B.P., and Thevissen, K. (2002). Plant defensins. Planta 216:193–202. 10.1007/s00425-002-0902-6.

van Zadelhoff, A., Meijvogel, L., Seelen, A.M., de Bruijn, W.J.C., and Vincken, J.P. (2022). Biomimetic Enzymatic Oxidative Coupling of Barley Phenolamides: Hydroxycinnamoylagmatines. J Agric Food Chem 70:16241–16252. 10.1021/acs.jafc.2c07457.

Visconti, S., Astolfi, M.L., Battistoni, A., and Ammendola, S. (2022). Impairment of the Zn/Cd detoxification systems affects the ability of. Front Microbiol 13:975725. 10.3389/fmicb.2022.975725.

Waters, B.M., Chu, H.H., Didonato, R.J., Roberts, L.A., Eisley, R.B., Lahner, B., Salt, D.E., and Walker, E.L. (2006). Mutations in Arabidopsis yellow stripe-like1 and yellow stripe-like3 reveal their roles in metal ion homeostasis and loading of metal ions in seeds. Plant Physiol 141:1446–1458. 10.1104/pp.106.082586.

Willmann, R., Lajunen, H.M., Erbs, G., Newman, M.A., Kolb, D., Tsuda, K., Katagiri, F., Fliegmann, J., Bono, J.J., Cullimore, J.V., et al. (2011). Arabidopsis lysin-motif proteins LYM1 LYM3 CERK1 mediate bacterial peptidoglycan sensing and immunity to bacterial infection. Proc Natl Acad Sci U S A 108:19824–19829. 10.1073/pnas.1112862108.

Xiao, F., Zhou, H., and Lin, H. (2025). Decoding small peptides: Regulators of plant growth and stress resilience. J Integr Plant Biol 67:596–631. 10.1111/jipb.13873.

Xue, R., Gao, N., Chen, J., Wu, Z., Sun, N., Li, Y., Gong, M., Zeng, R., Song, Y., Chen, D., et al. (2025). Hydroxycinnamic acid amides in rice: biosynthesis, distribution, function, and implication for crop development. Front Plant Sci 16:1525268. 10.3389/fpls.2025.1525268.

Yang, C., Liu, R., Pang, J., Ren, B., Zhou, H., Wang, G., Wang, E., and Liu, J. (2021). Poaceae-specific cell wall-derived oligosaccharides activate plant immunity via OsCERK1 during Magnaporthe oryzae infection in rice. Nat Commun 12:2178. 10.1038/s41467-021-22456-x.

Yu, X., Feng, B., He, P., and Shan, L. (2017). From Chaos to Harmony: Responses and Signaling upon Microbial Pattern Recognition. Annu Rev Phytopathol 55:109–137. 10.1146/annurev-phyto-080516-035649.

Zhou, Y., Zhou, B., Pache, L., Chang, M., Khodabakhshi, A.H., Tanaseichuk, O., Benner, C., and Chanda, S.K. (2019). Metascape provides a biologist-oriented resource for the analysis of systems-level datasets. Nat Commun 10:1523. 10.1038/s41467-019-09234-6.

