## Supplementary figures and images for "Molecular dissection of zinc-mediated immunity in Arabidopsis thaliana"

### Fig. S1

**Escudero FIGURE S1**

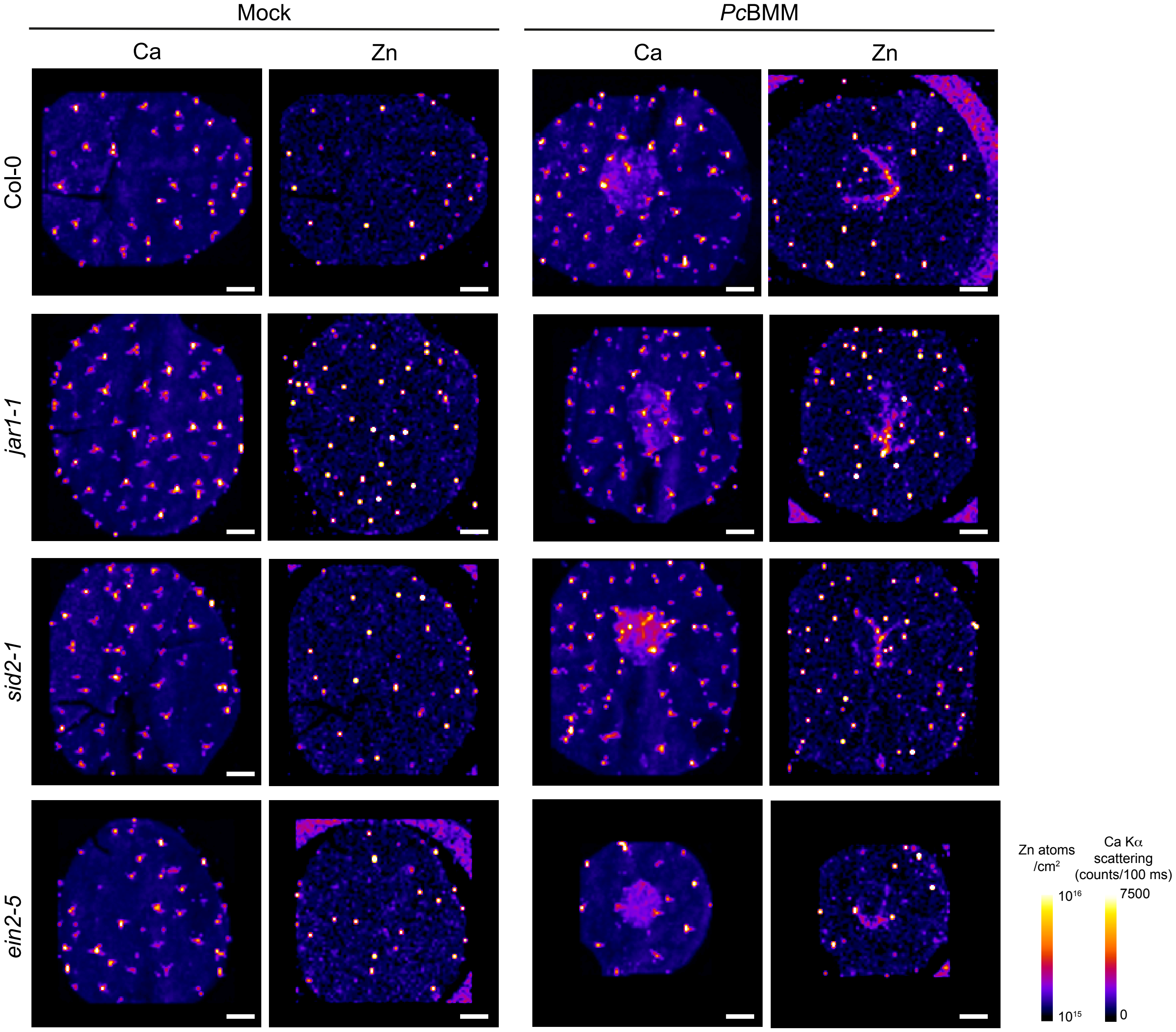
